# Atypical weighting of sensory evidence and priors in causality perception along the autism–schizotypy continuum

**DOI:** 10.1101/2025.06.11.659079

**Authors:** Gianluca Marsicano, David Melcher

**Affiliations:** Psychology Program, Division of Science, New York University Abu Dhabi, Abu Dhabi, United Arab Emirates; Center for Brain and Health, NYUAD Research Institute, New York University Abu Dhabi, Abu Dhabi, United Arab Emirates

## Abstract

The brain constructs a perceptual interpretation of the world by integrating sensory input with prior knowledge and expectations. Causality perception, a core example of this inferential process, enables observers to infer cause-effect relationships, such as whether a moving object causes another to move. Traditionally considered a low-level visual computation, causality perception is increasingly recognized as shaped by top-down dynamics, perceptual history, and individual predictive processing styles. Here, we examined how prior experience and cognitive-perceptual traits shape causal inference in 150 neurotypical individuals, using data-driven clustering to stratify participants along the autism–schizotypy (ASD–SSD) spectrum. Participants viewed dynamic collision events in which a moving circle contacted a stationary one, followed by variable temporal lags before the second object’s motion, and judged whether the interaction appeared causal or non-causal. Causality judgments were influenced by both physical timing (sensory-driven) and serial dependence on previous perceptual decisions (prior-driven). Hierarchical drift diffusion modeling (HDDM) revealed that SSD-like individuals showed a strong prior bias toward causality, greater serial dependence, and lower decision thresholds, reflecting a prior-dominated perceptual style. Conversely, ASD-like individuals exhibited reduced influence of perceptual history and higher decision thresholds, reflecting a conservative, data-driven style. Crucially, prior bias, serial dependence, and decision-making dynamics were strongly interrelated, revealing a hierarchical structure to perceptual inference across neurocognitive profiles. These findings demonstrate that causality perception emerges from predictive processes operating at multiple timescales and shaped by individual differences in neurocognitive style and perceptual flexibility.

**Significance Statement:** Accurate causality perception is a fundamental building block of cognition, supporting our ability to parse the sensory world, guide action, and construct coherent models of the environment. It provides a key example of perceptual inference, and thus how the brain integrates sensory input, prior expectations, and recent experience to shape subjective perception.

Here, we show that causality perception is not fixed, but varies systematically across individuals, mirroring broader variability in how different neurocognitive architectures balance sensory evidence and prior expectations along the autism–schizotypy (ASD–SSD) spectrum within the general population. By combining psychophysical data with hierarchical computational modeling, we reveal distinct perceptual inference styles along the autism–schizotypy continuum, ranging from sensory-driven to prior-driven processing.

These findings advance predictive processing accounts of individual variability and offer new insights into the mechanisms supporting flexible and adaptive perception in neurodiverse populations.

## Introduction

Human perception is not a passive reflection of sensory input but a dynamic inferential process through which the brain constructs coherent representations of the external world. A paradigmatic example is causality perception — the ability to perceive one event as the cause of another. This capacity is considered automatic and innate, present even in newborns [1], and plays a fundamental role in structuring the continuous stream of sensory input [2,3]. Critically, the ability to correctly perceive causality underlies core cognitive functions, shaping how we understand physical and social events, evaluate our own agency, and interpret events in terms of our beliefs in natural or supernatural forces at work [4].

A classic illustration of causality perception is the “*launching effect*” [5], where a moving object collides with a stationary one, causing the latter to appear set into motion. Critically, the impression of causality depends on timing: it emerges reliably when the second object moves within a narrow temporal window after the collision [5,6]. Even in the absence of actual physical contingency, sensory information can be integrated and experienced as causally related when presented within this temporal binding interval, whose breadth shows substantial interindividual variability [7].

Importantly, causality attribution extends beyond spatiotemporal cues, with individual differences in temporal binding thresholds reflecting the influence of top-down processes, including beliefs about physical principles [2], perceptual experience [8,9], and decision-making dynamics [10,11].

The predictive processing framework provides a powerful account of how causality perception emerges from the brain’s inferential dynamics. It posits that the brain operates as a hierarchical Bayesian inference engine, continuously generating predictions and updating them in light of sensory input [12]. In this framework, causality perception arises from the interplay between sensory evidence and top-down prior beliefs, with individuals varying along a continuum from sensory-driven to prior-driven processing styles [13,14].

These opposing styles become particularly salient in the contrast between autism spectrum disorder (ASD) and schizophrenia spectrum disorder (SSD). These conditions can be conceptualized as opposite poles on a continuum of predictive precision [14–18]. In ASD, perceptual inference is thought to rely heavily on sensory input and to underweight prior expectations [19]. Conversely, in SSD, inference is often described as overly prior-driven and characterized by rigid model updating [16]. However, both hypo- and hyperprior states can coexist within the same individual, with aberrant perception arising from misallocated precision across different levels of the predictive hierarchy [20]. Specifically, low-level sensory priors may be underweighted, leading to excessive and noisy prediction errors. To impose coherence on this uncertainty, the system may overcompensate by assigning excessive precision to high-level beliefs, as observed in SSD [20]. In line with this framework, core SSD features (e.g., delusions, ideas of reference, and anomalous perceptual experiences) have been linked to excessive reliance on internal models and widened temporal binding windows [21–23]. This imbalance fosters a stronger bias toward structured interpretations and faster evidence accumulation toward prior-consistent outcomes, manifesting as increased susceptibility to intentionality and causality attributions [9,24,25]. In contrast, individuals with ASD typically exhibit sensory-driven perceptual styles, characterized by greater reliance on sensory evidence and reduced top-down prior influence [19,26,27]. This style contributes to difficulties in integrating temporally distributed events into coherent causal narratives, as well as challenges with ambiguity resolution and contextual integration [19,28]. Overall, these diverging perceptual styles may reflect distinct precision-weighting mechanisms operating across hierarchical levels of inference, giving rise to unique neurocognitive profiles in both clinical and non-clinical populations [19,20].

Additionally, cause-effect interpretations depend on, and are continuously updated by, recent perceptual experience. This phenomenon, known as *serial dependence* [29], reflects the tendency for current perceptual judgments to be biased toward previous perceptual history (i.e., choice history bias; [30]), serving as an adaptive mechanism that promotes perceptual stability. Initially documented for low-level features, serial dependence also shapes higher-order processes such as causality perception: events following a causal impression are more likely to be perceived as causal [8]. However, the strength of serial dependence varies across individuals. When overly strong, it may foster perceptual rigidity; when too weak, it may impair the adaptive updating of internal models [31,32]. This opposing influence of perceptual history remains evident along the ASD-SSD continuum. SSD traits are associated with excessive reliance on perceptual history, which can reinforce biased causal attributions [9], whereas ASD profiles are linked to a reduced influence of recent experience, resulting in slower updating of internal representations [31,33,34].

Taken together, these findings highlight that causality perception arises from the interplay of sensory processing, top-down mechanisms, and the dynamic updating of internal representations, components that are dynamically weighted and may interact differently across neurocognitive phenotypes [16,20,21,28]. While these components are often investigated separately, they do not operate in isolation; rather, each stage of the perceptual inference process dynamically interacts with the others to shape the final perceptual outcome.

In the current study, we aimed to disentangle how sensory input and top-down components of perceptual inference interact to shape causality perception in the neurotypical population. Building on previous evidence, we hypothesized diverging predictive styles in causality inference, consistent with hierarchical models in which perception emerges from the precision-weighted integration of bottom-up sensory signals and top-down expectations [20]. We applied this framework dimensionally, reasoning that individual differences in cognitive-perceptual traits may reflect variability in how perceptual precision is distributed across hierarchical levels of sensory evidence and prior expectations, influencing how causal events are perceived. Based on this rationale, we expected that individuals with SSD-like profiles, marked by cognitive-perceptual atypicalities such as delusions and hallucinations, would exhibit increased causality perception, reflecting a prior-dominated inference style in which perceptual history reinforces biased causal attributions [9,16,35]. In contrast, individuals with ASD-like profiles were expected to rely more strongly on sensory evidence, resulting in slower model updating and a reduced influence of recent perceptual history [28,31,34,36,37]. Finally, we expected individuals without prominent cognitive-perceptual atypicalities to exhibit a more balanced predictive style, flexibly integrating sensory input, prior models, and perceptual history.

To test these hypotheses, we used a *Michotte launching paradigm* (Fig. 1), in which 150 neurotypical participants judged the perceived causality of collision events with varying temporal lags. We characterized phenotypic variability along the ASD-SSD continuum by applying a data-driven k-means clustering approach to self-reported perceptual, cognitive, socio-affective, and communicative traits, stratifying participants into three subgroups: ASD-like, SSD-like, and low-trait (LT) profiles. We then examined how these individual differences influenced causality perception, first by assessing temporal integration thresholds for causality (*point of subjective causality*, PSC) and evaluating the extent to which prior perceptual history (serial dependence) biased current causal judgments. Finally, we applied a computational Hierarchical Drift Diffusion Model (HDDM; [38]) to decompose perceptual decision-making and characterize how starting bias, evidence accumulation, and decision thresholds varied across profiles. Together, this approach enabled us to disentangle the sensory, top-down, and history-dependent components of perceptual inference in causality perception.

**Figure 1.**
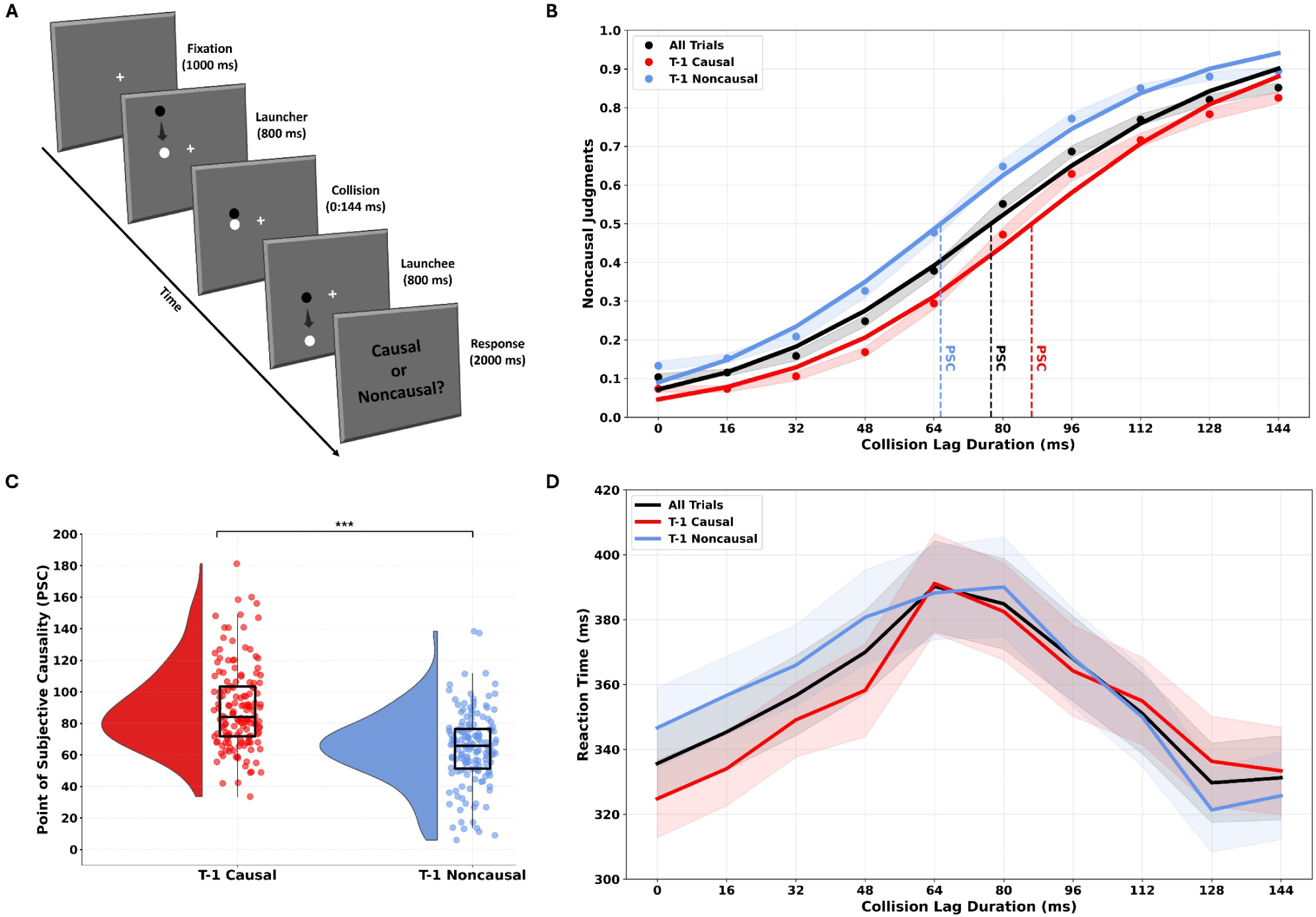
Experimental design and overall results across participants. A) Each trial began with a central fixation cross (1 s), followed by the appearance of two circles on one side of the screen. The first circle moved for 800 ms toward a stationary target at midline and collided with it. To manipulate causality perception, the onset of motion in the second circle was delayed by one of nine temporal lags (0-144 ms, in 17 ms steps). After the motion sequence, participants made a forced-choice judgment (“Causal or Noncausal?”). B) Mean non-causal response rate across collision time lags for all participants. Psychometric functions are plotted for all trials (black), trials following a causal response (red), and trials following a non-causal response (blue). Vertical dotted lines represent the Point of Subjective Causality (PSC; 50% threshold) for each condition. Shaded areas indicate the standard error of the mean (SEM). C) Raincloud plot illustrating the PSC difference between trials preceded by a causal (blue) versus non-causal (red) judgment. PSCs were significantly lower following causal trials, indicating a serial dependence effect. Half-violin plots display the distribution, bars represent the mean, and individual dots show participant-level data; error bars reflect SEM. D) Mean reaction times (RTs) at each collision lag, shown separately for all trials (black), t-1 causal trials (red), and t-1 non-causal trials (blue). RTs peaked at intermediate lags, where causality judgments tend to be most ambiguous, and were shorter at clear-cut lag intervals. Shaded areas represent SEM.

## Results

### Cluster Analysis: Individual Profiles Along the ASD-SSD Continuum

To investigate how individual differences influence the construction of causality perception, we first characterized the multiple potential phenotypes observable in our sample (n=150 neurotypical participants) computing a data-driven k-mean clustering approach on self-reported perceptual, cognitive, socio-affective and communicative styles derived from the Autism Quotient (AQ; [39]) and Schizotypal Personality Questionnaire (SPQ; [40]). Silhouette scores of the clusters for values of *K* ranging from 1 to 10 were calculated to determine the optimal number of clusters (*K*), selecting the *K* producing the highest silhouette score. According to the silhouette coefficient (0.36), the optimal clustering solution was 3 *K*, suggesting a reasonable 3-clusters structure ([41]; see Table 1 for clustering summary).

**Table 1.** Cluster solutions derived from the clustering analysis. The table presents R², Bayesian Information Criterion (BIC), and Silhouette scores for the cluster solutions obtained. The analysis revealed higher R² and Silhouette score for the 3-cluster structure stratifying participants in ASD-like traits, Low Traits, and SSD-like Traits.

| N Clusters | R<sup>2</sup> | BIC | Silhouette |
| --- | --- | --- | --- |
| 2 | 0.290 | 1620.39 | 0.28 |
| 3 | 0.532 | 1143.29 | 0.36 |
| 4 | 0.520 | 1134.16 | 0.34 |
| 5 | 0.420 | 1187.39 | 0.31 |

In line with previous evidence [14,42], the 3-cluster structure obtained in our analyses (Fig. 2) comprised: 1) a first cluster (n=44, 29.3% of participants), named “ASD-like”, composed with individuals displaying high scores on the socio-affective dimension (SPQ subscales: *no close friends, social anxiety, constricted affect*; AQ subscales: *social skills, communication, attentional to detail, attention switching, imagination*), and low scores in the cognitive-perceptual domain (*SPQ: ideas of reference, odd beliefs/magical thinking, unusual perceptual experiences, odd behaviour, odd speech, suspiciousness*); 2) a second cluster (n=50, 33.3% of participants), called “SSD-like” was characterized by individuals showing high scores on the cognitive-perceptual dimension (SPQ subscales: *ideas of reference, odd beliefs/magical thinking, unusual perceptual experiences odd behaviour, odd speech, suspiciousness;* AQ subscales: *attentional switching*), and low scores in the socio-affective dimension (SPQ subscales: *no close friends, social anxiety, constriced affect*; AQ subscales: *social skills, communication, attention to detail, communication, imagination*); 3) a third cluster (n=56, 37.3% of participants), called “Low Traits” (LT), composed of individuals with low scores in both AQ and SPQ subscales. The subscales *attentional switching* and *imagination* (AQ) showed comparable moderate scores both on ASD-like and SSD-like clusters, and lower scores in LT cluster. Overall, these results align with previous findings highlighting a diametric relationship between autistic and schizotypal traits, representing opposing poles of a personological continuum among individuals exhibiting atypicalities in socio-communicative-affective and cognitive-perceptual dimensions [14].

**Figure 2.**
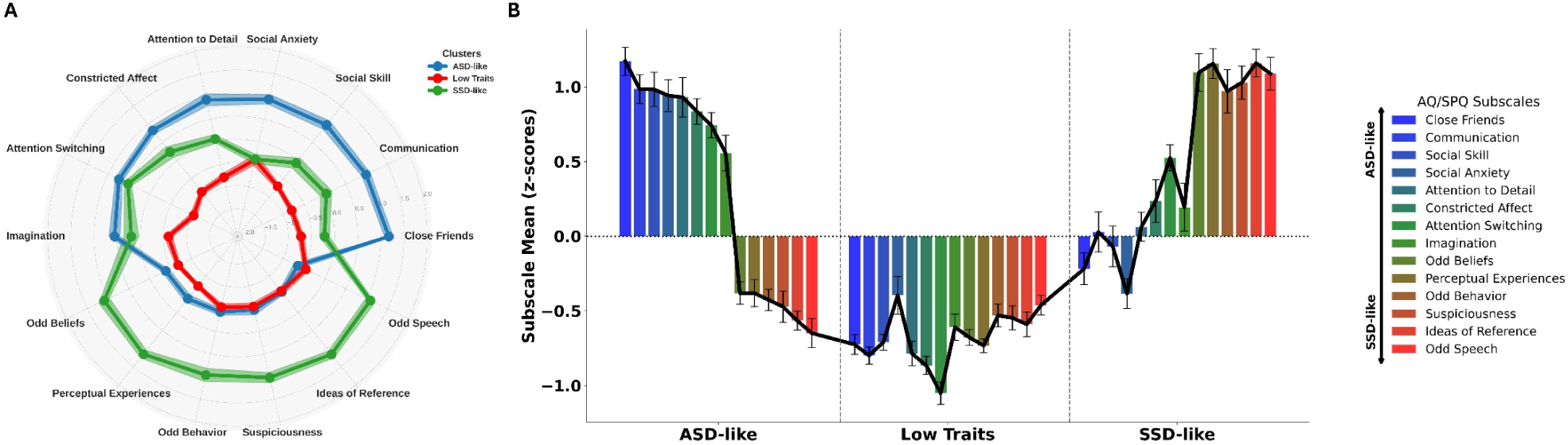
Data-driven k-mean cluster analysis: ASD-SSD like sub-phenotypes. The results highlight a diametric relationship between schizotypal and autistic traits, representing opposing poles of a continuum among individuals exhibiting atypicalities in cognitive-perceptual and socio-communicative-affective dimensions. Panel A (radar plot) and Panel B (bar plot) depict the mean z-scores for each subscale of the SPQ and AQ questionnaires across the ASD-like, Low Traits, and SSD-like clusters. For each cluster, the plots highlight the contribution of subscales within each cluster, revealing high socio-communicative-affective atypicalities in ASD-like cluster (SPQ subscales: no close friends, social anxiety, constricted affect; AQ subscales: social skills, communication, attentional to detail, attention switching, imagination), high cognitive-perceptual anomalies in SSD-like cluster (ideas of reference, odd beliefs/magical thinking, unusual perceptual experiences odd behaviour, odd speech, suspiciousness; AQ subscales: attentional switching), and low scores across subscales for Low Traits cluster. The error bars indicate the standard error of the mean (SEM).

### Atypically Increased Causality Inference in SSD-like Profiles

We first examined whether judgments of the causality of collision events during *Michotte launching paradigm* (Fig. 1; [5]) varied across different collision lags as a function of participant clusters identified through k-means cluster analysis, indexing individuals Point of Subjective Causality (PSC) via logistic modelling of non-causal response rate. Accordingly, we performed a one-way ANOVA on PSC with cluster as between-subjects factor (three-levels: ASD-like, Low Traits, SSD-like). The results revealed a significant main effect of Cluster (F_(2,147)_ = 6.351, *p* = 0.002, η2p = 0.08), suggesting distinct profiles of causality perception within our sample of participants. SSD-like participants showed heightened PSC (M = 88.62, SD = 26.25), as compared to ASD-like (M = 74.23, SD = 22.8; *p* = .01, Cohen’s *d* = 0.589) and Low Traits (M = 73.08, SD = 23.92; *p* = .004, Cohen’s *d* = 0.637) groups. On the contrary, no significant difference emerged from the comparison between ASD-like and Low Traits (*p* = .814, Cohen’s *d* = -0.047). Overall, these findings offer novel insights into variability within the neurotypical population, consistent with previous evidence highlighting heightened causality perception in individuals showing prominent cognitive-perceptual atypicalities (e.g., delusional ideation, ideas of reference; e.g., [43]).

### Dependence on Perceptual History: Attenuated in ASD-like, Amplified in SSD-like participants

Subsequently, we investigated potential individual differences in the extent to which prior perceptual history influenced current causality perception by examining how preceding responses affected the current causality judgments across the different subgroups (i.e., *serial dependence*; trial n-1). Previous perceptual experience is typically integrated with current sensory information to support perceptual continuity over time (Fischer & Whitney, 2014). However, the effect of prior perceptual experience often shows substantial interindividual variability, particularly along the ASD-SSD spectrum. Individuals with high ASD traits tend to rely less on prior experience, favoring current sensory input [36], whereas individuals with SSD traits often exhibit a stronger reliance on stable prior models [14], reflecting distinct differences in how past information is integrated into current perceptual decisions.

Accordingly, PSC was separately indexed for trials following causal (t-1 causal) and non-causal (t-1 non-causal) judgments. A 2×3 repeated-measures ANOVA was then conducted, with t-1 judgment (two-levels: t-1 causal, t-1 non-causal) as the within-subjects factor and cluster (three-levels: ASD-like, Low Traits, SSD-like) as the between-subjects factor. Consistently with previous findings [8], perceptual history significantly biased causality judgments (F_(1,147)_ = 158.335, *p* < 0.001, η2p = 0.519; Fig. 1), with causal responses increased following causal trials (M = 89.74, SD = 26.37) as compared to non-causal judgments (M = 64.39, SD = 23.24; *p* < .001, Cohen’s *d* = 1.039).

Importantly, such analysis revealed a significant interaction between t-1 judgment and Cluster factors (F_(2,147)_ = 10.115, *p* < 0.001, η2p = 0.121; Fig. 3). Within-group post-hoc comparisons revealed that prior perceptual history significantly influenced current causality judgments across all clusters, with an increased likelihood of causal responses following causal trials, and a corresponding increase in non-causal responses following non-causal trials (ASD-like = t–1 causal: M = 82.03, SD = 21.51; t– 1 non-causal: M = 66.76, SD = 26.36; *p* < .001, Cohen’s *d* = 0.637; Low Traits = t–1 causal: M = 84.32, SD = 19.94; t–1 non-causal: M = 61.77, SD = 23.14; *p* < .001, Cohen’s *d* = 0.935; SSD-like = t–1 causal: M = 102.58, SD = 31.74; t–1 non-causal: M = 65.32, SD = 20.46; *p* < .001, Cohen’s *d* = 1.545; Fig. 3). On the other hand, consistent with the notion of heightened causality perception in SSD, between-group post hoc comparisons showed that SSD-like participants exhibited significantly higher PSCs and causality judgments following causal trials compared to both ASD-like (*p* < .001, Cohen’s *d* = 0.852) and Low Traits (*p* < .001, Cohen’s *d* = 0.757) groups, while no significant differences were found in the remaining comparisons (all *p*-values > .58).

**Figure 3.**
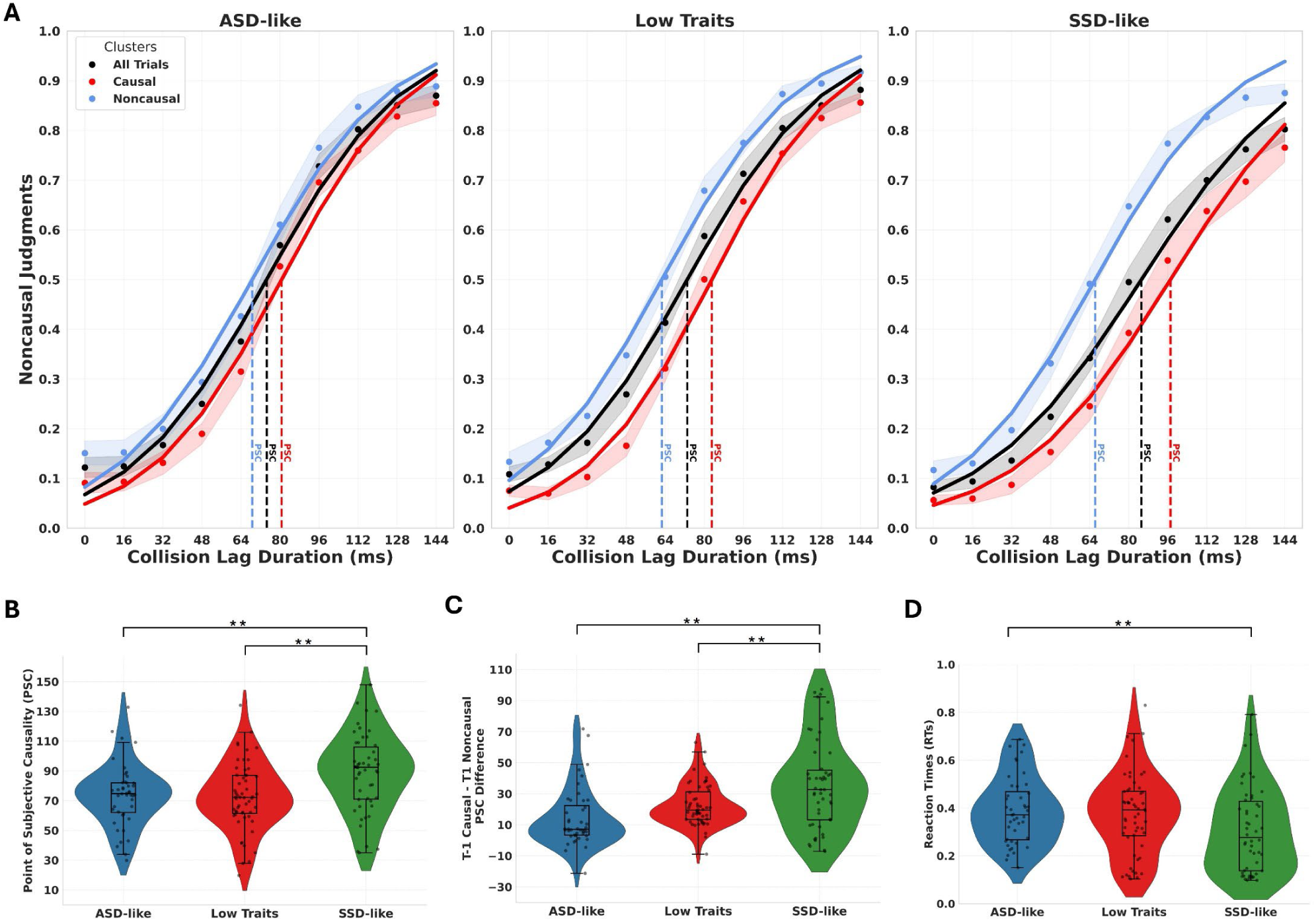
Group-level differences in causality inference. A) Psychometric functions showing the proportion of non-causal judgments as a function of collision time lag for each subgroup (left panel: ASD-like; central panel: Low Traits; right panel: SSD-like). Curves are plotted for all trials (black), trials following causal responses (red), and trials following non-causal responses (blue). Vertical dashed lines indicate the Point of Subjective Causality (PSC; 50% threshold) for each condition. Shaded areas represent standard error of the mean (SEM). B) Violin plots of PSC values across subgroups. SSD-like participants showed significantly higher PSCs compared to both ASD-like and Low Traits groups, indicating a greater temporal tolerance for inferring causality and increased causality judgments. C) PSC difference scores (t-1 causal vs. t-1 non-causal) indexing serial dependence across subgroups. SSD-like participants exhibited significantly greater serial dependence effects than both ASD-like and Low Traits groups, suggesting a stronger influence of recent perceptual history on causal judgments. D) Mean reaction times (RTs) across groups. SSD-like participants responded faster than ASD-like and Low Traits individuals, suggesting reduced decisional caution and a stronger prior-driven response style. In all plots, boxplots represent the median and interquartile range; violins depict the distribution; individual dots reflect participant-level data. Asterisks indicate significance levels (*=p<.05; **=p<.01; ***=p<.001).

To further assess the extent to which prior perceptual history influenced current causality judgments across clusters, we computed the PSC difference between *t-1* non-causal and *t-1* causal trials, with larger differences indicating a stronger serial dependence effect. We then conducted a one-way ANOVA on the resulting PSC difference index, with cluster as the between-subjects factor (three-levels: ASD-like, Low Traits, SSD-like). As expected, the results revealed a significant main effect of Cluster (F_(2,147)_ = 10.115, *p* < 0.001, η2p = 0.121; Fig. 3), with higher PSC difference in the SSD-like group (M = 37.25, SD = 34.45) with respect to ASD-like (M = 15.36, SD = 24.16; *p* < .001, Cohen’s *d* = 0.902) and Low Traits (M = 22.55, SD = 13.4; *p* = .004, Cohen’s *d* = 0.606) groups. Overall, this pattern of results highlights distinct individual profiles in the weighting of prior perceptual history in causality judgments, with SSD-like individuals showing a stronger influence of previous perceptual experience, whereas ASD-like participants showed a reduced sensitivity to perceptual history.

### Diverging Reaction Times in Causality Judgment Along the ASD-SSD Continuum

Next, we explored potential interindividual differences in the temporal dynamics of causal judgments by comparing Reaction Times (RTs) at each collision lag across clusters. Alongside accuracy, fast RTs can indicate efficient processing of cause-effect relationships, as atypicalities in response speed may reflect disruptions in the perception of temporal contiguity [11].

Accordingly, we first conducted a 10×3 repeated-measures ANOVA (rmANOVA) on RTs across all trials to assess whether the speed of causality judgments was modulated by collision lag (within-subject: 10 levels) and cluster (between-subjects: ASD-like, Low Traits, SSD-like). The results revealed a main effect of cluster (F_(2,147)_ = 3.43, *p* = 0.035, η2p = 0.045; Fig. 3D), with SSD-like participants (M = .307 seconds, SD = .177 seconds) exhibiting faster RTs compared to ASD-like participants (M = .388 seconds, SD = .14 seconds; *p* = .048, Cohen’s *d* = 0.55), while the remaining between-group comparisons were not statistically significant (all *p*-values > .088).

Interestingly, regardless of the between-subjects factor Cluster, such analysis revealed a main effect of collision lag (F_(5.16,759.53)_ = 13.071, *p* < 0.001, η2p = 0.082; Fig. 1D). Post hoc comparisons revealed that at intermediate collision time lags, participants exhibited significantly slower RTs compared to short and long-time lags, where the perception of causality tends to be more clear-cut (see Fig. 1D). However, the rmANOVA performed on RTs did not reveal a significant interaction between Cluster and Time Lag (F_(18,1323)_ = 1.125, *p* = 0.32, η2p = 0.015).

Overall, these findings suggest that both individual profiles and collision timing significantly influence the temporal dynamics of causal perception. Specifically, SSD-like participants exhibited faster reaction times compared to the ASD-like group, aligning with previous findings that individuals with elevated SSD-like traits often prioritize speed over accuracy [10,25]. This pattern may reflect a cognitive bias toward perceiving collisions as causal events, potentially driven by a more inflexible decision-making style that favors rapid judgments regardless of perceptual ambiguity.

### Modelling Perceptual Decision-making Dynamics Along the ASD-SSD Continuum

Upon the observed differences across groups, we next computed a Hierarchical Drift Diffusion Model (HDDM; [38]) to gain deeper insight into the underlying decision-making mechanisms shaping causality judgments. In this realm, HDDM offers a powerful Bayesian computational framework for capturing individual differences in sensory evidence evaluation and decision-making response strategies [38,56]. Accordingly, here we fitted HDDM to the RTs distributions for causality responses (see: *Methods* and *Supplementary Materials*), separately for each cluster, to investigate whether individual differences in visual causality judgments may be explained by variations in the starting point of the accumulation process (initial *bias* in favor of one option), the speed at which evidence is accumulated (*drift rate*), or the amount of information required to make a decision (*decision boundary*). Next, we characterized the posterior distribution of each parameter by indexing its median value and high-density interval (HDI), and compared between-group posterior distributions by subtracting HDIs; if the HDI distribution of differences did not include zero, the condition effect was considered credible and thus statistically significant ([45], see: Methods).

Consistent with the hypothesis of heightened prior beliefs in cause-effect attribution, the SSD-like group exhibited a stronger initial bias (*z*) toward causality compared to both the Low Traits group (HDI_difference_ = [0.057; 0.018]) and the ASD-like group (HDI_difference_ = [0.006; 0.054]; Fig. 4A). Conversely, no credible differences emerged between the ASD-like and Low Traits groups (HDI_difference_ = [-0.028; 0.018]; Fig. 4A). Importantly, SSD-like individuals not only initiate decision-making with a stronger initial bias, but also accumulated evidence towards the causal response more rapidly (Fig. 4B). The SSD-like group showed a credible increased drift rate (*v*) toward causal responses with respect to Low Traits participants (HDI_difference_ = [-0.31; -0.011]). On the other hand, no credible differences emerged from the comparison between the SSD-like and ASD-like groups (HDI_difference_ = [-0.014; 0.027]), nor between the ASD-like and Low Traits (HDI_difference_ = [-0.013; 0.023]). Additionally, decision thresholds (*a*) were reached at different times across groups (Fig 4C). SSD-like individuals made causality judgments more rapidly and with lower decision boundaries compared to both the Low Traits (HDI_difference_ = [–0.229, –0.027]) and ASD-like groups (HDI_difference_ = [–0.283, –0.07]), suggesting a more impulsive decision-making policy in SSD-like participants, and a more cautious response strategy in the ASD-like group.

**Figure 4.**
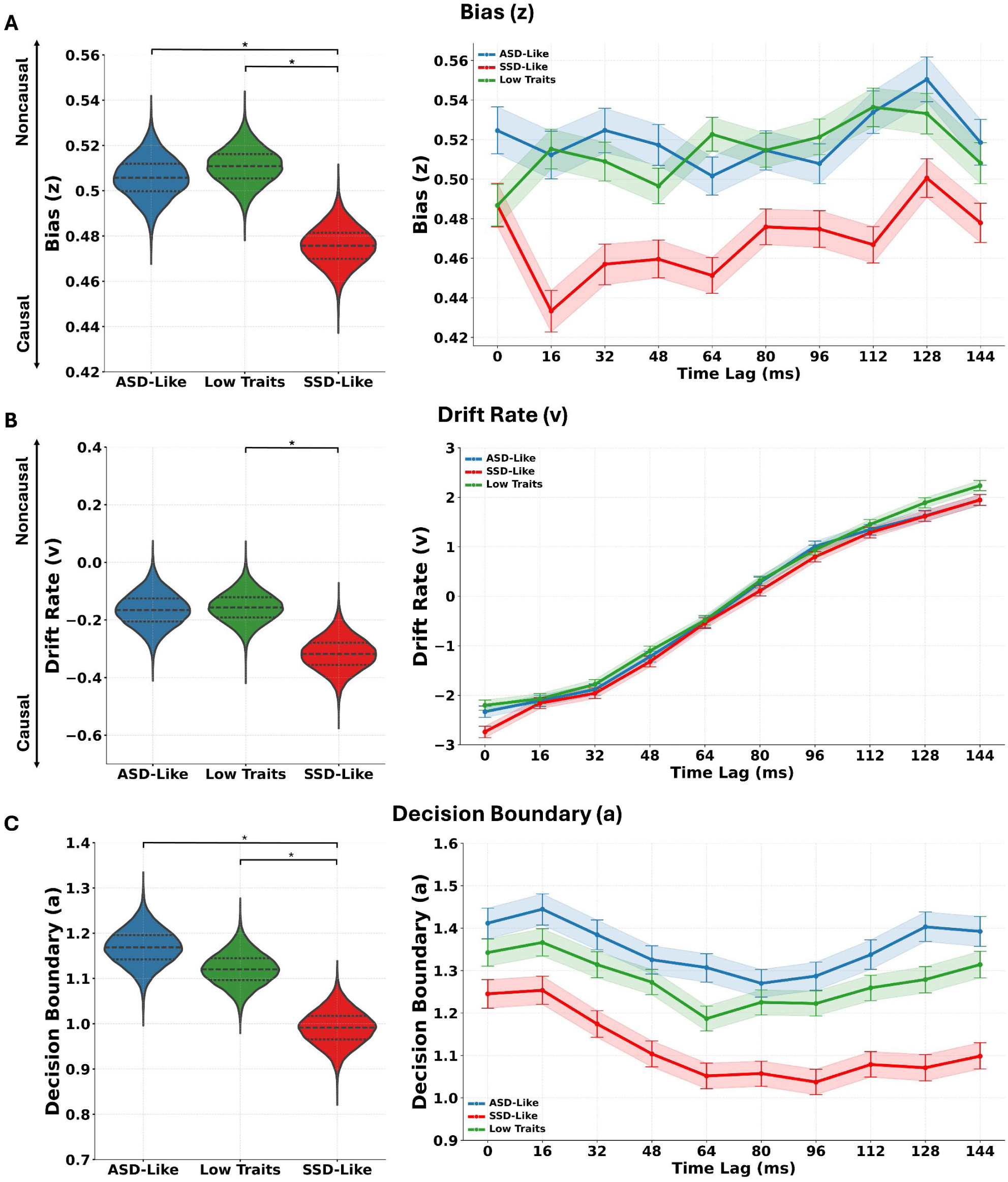
Hierarchical Drift Diffusion Model: dissociable perceptual decision-making profiles along the ASD-SSD continuum. A) Violin plots (left) and time course plots (right) of starting bias (z) toward causal responses across groups. SSD-like participants exhibited a significantly stronger initial bias toward causality compared to both Low Traits and ASD-like groups, consistent with a prior-driven inferential style. No credible difference emerged between ASD-like and Low Traits groups. B) Drift rate (v), reflecting the speed of evidence accumulation toward a decision. SSD-like individuals showed a significantly higher drift rate for causal responses relative to Low Traits participants, indicating faster commitment to causal judgments. Drift dynamics across time lags (right) revealed a consistent pattern across groups: evidence accumulation was fastest at short lags (strong causal impression), slowed near perceptual ambiguity (intermediate lags), and reversed at longer lags (favoring non-causal judgments). C) Decision boundary (a), indexing the amount of evidence required before making a decision. SSD-like participants showed lower decision thresholds compared to both ASD-like and Low Traits groups, suggesting a more impulsive decision policy. In contrast, ASD-like individuals exhibited the highest boundaries, reflecting greater decisional caution. Shaded areas in time course plots represent SEM. Asterisks denote credible differences based on HDI comparisons.

Additionally, we explored potential interactions between clusters and collision lags for each HDDM parameter (Fig. 4; see: *Methods*). Overall, the between-group credible differences previously identified for each parameter remained consistent across time lags. Across all groups, evidence accumulation was faster toward causality responses (drift rate) at short time lags, minimal at intermediate lags where perceptual ambiguity was highest, and faster toward non-causal judgments at long time lags (Fig. 4B). Regarding decision boundaries, higher thresholds were observed at shorter time lags across groups, suggesting a more cautious response strategy compared to longer time lags (Fig. 4C).

Overall, the HDDM results revealed distinct decision-making profiles underlying causality judgments: SSD-like individuals exhibited a stronger initial bias toward causality, faster evidence accumulation, and lower decision boundaries, whereas ASD-like individuals tended toward a more cautious and conservative decision-making strategy.

### Decision-Making Dynamics Interact with Perceptual History to Shape Causal Inference

Finally, we performed Pearson correlation analyses to test whether individual differences in decision dynamics predict variability in PSC and serial dependence from perceptual history.

Regarding PSC, it was negatively correlated with the starting bias (z; *r* = –0.68, *p* < 0.001; Fig. 5A) and drift rate (v; *r* = –0.88, *p* < 0.001; Fig. 5B), but not with decision boundary (a; *r* = –0.12, *p* = 0.11; Fig. 5A). This pattern of results suggests that stronger priors (i.e., starting bias) toward causal interpretations reduce sensitivity in detecting accurate cause-effect relationships in colliding visual stimuli, as reflected in higher PSC values. Additionally, the speed of evidence accumulation (i.e., drift rate) was faster for causal judgments in participants with lower PSCs, and faster for non-causal judgments in those with higher PSCs, indicating that drift dynamics align with individual differences in causality perception.

**Figure 5.**
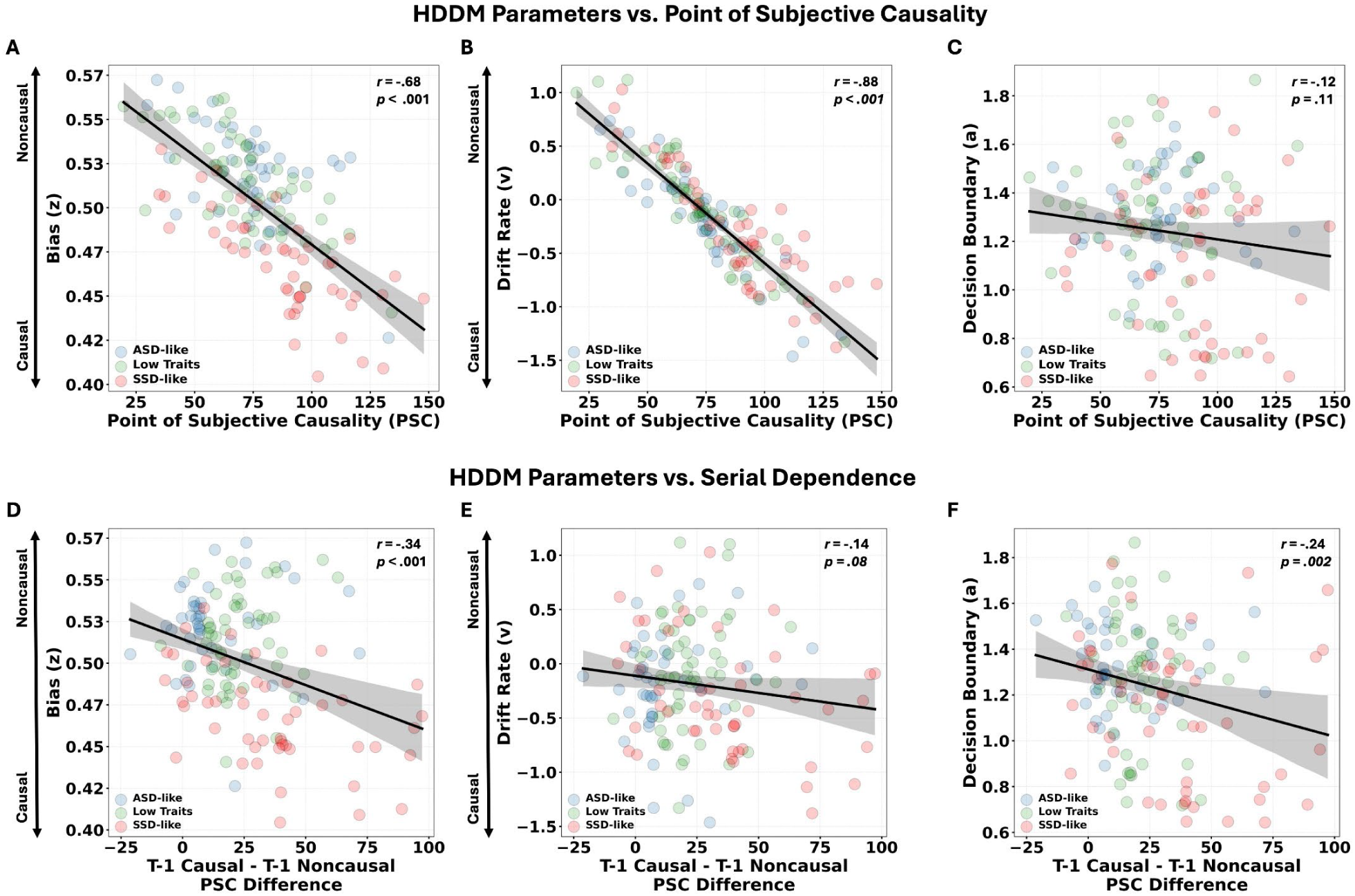
Causality Perception emerges from the interaction of perceptual priors and decision dynamics. A-C) Scatterplots depicting Pearson correlations between HDDM parameters and PSC across all participants. Strong negative associations were observed between PSC and starting bias (A; *r* = –.68, p < .001) and drift rate (B; *r* = –.88, p < .001), indicating that individuals with a stronger initial bias and faster accumulation of evidence toward causal interpretations (i.e., lower *z* and *v* values) required longer temporal delays to perceive non-causal contingencies (i.e., increased causality judgments). No significant association emerged between PSC and decision boundary (C; *r* = –.12, p = .11). D-F) Correlations between HDDM parameters and serial dependence effect (i.e., difference in PSC following t-1 causal vs. t-1 non-causal responses). Greater serial dependence was associated with stronger starting bias toward causality (D; *r* = – .34, p < .001) and lower decision boundaries (F; *r* = –.24, p = .002), suggesting that individuals with stronger priors and reduced decisional caution were more influenced by recent perceptual history. No significant relationship was found between serial dependence and drift rate (E; *r* = –.14, p = .08). Together, these findings indicate that stable internal priors (starting bias), dynamic perceptual updating (serial dependence), and decisional strategies (decision boundary) jointly shape causality perception, suggesting a hierarchical inferential process differently modulated across individual profiles along the ASD-SSD continuum. Shaded bands represent 95% confidence intervals.

On the other hand, a stronger serial dependence on previous perceptual history (i.e., difference between t-1 causal vs. t-1 non-causal trials) was negatively correlated with the starting bias (z; r = – 0.34, p < 0.001; Fig. 5A) and decision boundary (a; r = –0.34, p = 0.002; Fig. 5B), but not with drift rate (v; r = –0.14, p = 0.08; Fig. 5A). First, these results highlight that perceptual history and initial bias, although reflecting different sources of influence and operating at different timescales, interact strongly in reinforcing models of causality attribution. Thus, it is conceivable that strong serial influence from prior trials may reinforce the initial bias by stabilizing causal interpretations across time through a perceptual feedback loop in which biased perceptual decisions become increasingly self-confirming. Furthermore, higher decision thresholds predicted reduced dependence on perceptual history in detecting cause-effect relationships, suggesting that participants who rely more on sensory evidence on a trial-by-trial basis may adopt a more cautious decision-making strategy, requiring more time to reach a decision.

## Discussion

In the current study, we sought to characterize and interpret the marked individual differences that shape causality perception. Our findings support the emerging view that causality perception is not a fixed, bottom-up process, but rather a multi-stage hierarchical inferential process. Across all participants, judgments of causality were modulated by both the temporal structure of the stimulus [5,11], and by recent perceptual experience [19], which introduced systematic serial dependence into perceptual decisions, reflecting an attractive mechanism that stabilizes perceptual inference over time [8]. Moreover, individual differences in these processes were strongly linked to variability in top-down predictive models [46,47]). This pattern supports contemporary frameworks in which perceptual inference arises from the interaction of bottom-up sensory input, prior beliefs, and dynamic updating mechanisms [8,9,20]. Critically, these inferential dynamics did not operate uniformly across participants. Rather, they reflected systematic trait-dependent variability in how priors and sensory evidence are weighted, updated, and integrated in coherent perceptual representations [14,35].

### Characterizing Individual Profiles along the ASD-SSD continuum

A central question motivating our study was whether trait-dependent differences in cognition and perception influence how cause-effect attributions are constructed. To capture this interindividual variability, we adopted a data-driven clustering approach, applying k-means clustering to self-reported perceptual, cognitive, socio-affective, and communicative styles. This strategy allowed us to identify naturally occurring phenotypic profiles, rather than imposing arbitrary thresholds. We identified three clusters: participants with high autistic traits (ASD-like), participants with high schizotypal traits (SSD-like), and participants with low traits (LT) in both dimensions.

Our clustering structure aligned with prior evidence suggesting a diametric relationship between ASD- and SSD-like traits in the general population, whereby socio-communicative atypicalities are often accompanied by intact cognitive-perceptual abilities, and cognitive-perceptual atypicalities may coexist with relatively preserved social-affective functioning [14,15,18,22].

Importantly, these profiles are also thought to reflect opposing poles on a continuum of predictive precision [16,48], with ASD profiles exhibiting sensory-driven inference [19], and SSD profiles exhibiting over-weighted priors and inflexible model updating [16,17,20]. Accordingly, our stratification thus provided an ideal framework to test whether these opposing inference styles would manifest in systematic differences in causality perception, perceptual decision dynamics, and history-dependent updating.

### Perceiving Cause in Time: Divergent Inference Styles Along the ASD-SSD Continuum

First, our results showed that SSD-like participants exhibited heightened causality perception, as reflected in higher temporal integration thresholds (i.e., point of subjective causality) compared to both LT and ASD-like participants. This finding aligns with prior work linking cognitive-perceptual atypicalities (e.g., delusional ideation, ideas of reference) to increased cause-effect attribution even in presence of strong sensory cues [9]. This imbalance may result in perceptual experiences that are coherent but inaccurate, ranging from hallucinations and delusions in clinical populations to subtler misattributions in subclinical manifestations. Additionally, core SSD-like features have been consistently associated with widened temporal binding windows for visual events [21–23], contributing to distorted sensory processing. This imbalance may explain the stronger bias toward structured interpretations, manifesting as increased susceptibility to temporal binding and causality attributions [9,23]. These findings are consistent with recent hierarchical models of perceptual inference, suggesting that such biases may stem from misallocated precision across levels of the perceptual hierarchy [20]. Specifically, underweighted low-level sensory priors may result in increased prediction error, prompting the system to over-rely on high-level beliefs to impose perceptual coherence, ultimately driving rigid causal interpretations, even under strong sensory evidence.

Interestingly, individual profiles also significantly predicted the temporal dynamics of causal perception, suggesting that differences in perceptual inference extend beyond accuracy alone. SSD-like participants exhibited faster response times, potentially reflecting a rigid decision-making style in which strong priors drive rapid, confident judgments even under sensory uncertainty [10,25]. Conversely, ASD-like participants exhibited slower response times, consistent with a more sensory-driven, cautious approach to perceptual decisions. This pattern aligns with evidence that ASD traits are associated with reduced reliance on priors, enhanced sensory precision, and greater detail-focused processing, resulting in a difficulty to integrate temporally distributed events into coherent narratives [19,26,28]. Together, these findings suggest that predictive processing differences along the ASD-SSD spectrum influence not only what is perceived, but also how perceptual decisions unfold over time.

### Priors and Perceptual Decision-Making Dynamics along the ASD-SSD continuum

Differences in causality perception across the ASD–SSD continuum emerged both behaviorally and computationally, as revealed by Hierarchical Drift Diffusion Modeling (HDDM). Consistent with our hypotheses, the SSD-like group exhibited a stronger initial bias (*z*) toward causality compared to both the Low Traits and ASD-like groups. This bias was complemented by a faster rate of evidence accumulation toward causal responses, suggesting that SSD-like individuals not only begin the decision process with stronger expectations of causality but also arrive at those judgments more swiftly. Additionally, the SSD-like group demonstrated lower decision thresholds, indicating a reduced need for sensory evidence before committing to a decision, aligning with prior findings showing that individuals within the SSD spectrum often favor speed over deliberation and show diminished tolerance for perceptual ambiguity [10,25]. Thus, our findings point to an overarching perceptual style in SSD-like individuals, wherein causal structure is readily inferred, even in the presence of strong temporal cues, consistent with predictive processing accounts of psychosis spectrum [13,20,35]. In contrast, ASD-like participants did not show atypical prior bias or drift rate, but exhibited higher decision thresholds, suggesting a more conservative decision policy, requiring greater sensory evidence before committing to a perceptual judgment. This computational profile aligns with sensory-driven inference models of ASD, in which increased sensory weighting and reduced prior influence promote greater deliberation and caution in perceptual decisions [28,49]. Taken together, these computational findings strongly support the existence of distinct perceptual inference profiles underlying causality judgments, providing a characterization of distinct precision-weighting strategies across hierarchical levels.

### Not Just Stable Priors: Perceptual History as a Window into Predictive Processing

Beyond stable priors, our findings show that causality judgments are also shaped by dynamic influences from recent perceptual experience (i.e., *serial dependence*). We observed robust serial dependence effects across all groups, whereby prior judgments (*trial n-1*) significantly biased current causality perception. Crucially, the influence of perceptual history varied systematically across profiles, with SSD-like individuals exhibiting the strongest serial dependence effect and ASD-like individuals the weakest. These findings align with previous evidence suggesting that individuals with high SSD-like atypicalities place greater weight on prior perceptual experiences, reinforcing biased causal interpretations through a rigid, prior-consistent inferential style [9,35]. Additionally, this pattern is consistent with hierarchical predictive models of SSD, in which underweighted low-level sensory priors increase reliance on recent high-level predictions to stabilize perception, fostering perceptual rigidity [20]. In contrast, the reduced susceptibility to perceptual history observed in ASD-like individuals may reflect an absent or slower updating of internal representations and a more sensory-driven perceptual style, as often reported in the ASD spectrum [13,31,37]. Together, these findings suggest that differences in how priors are dynamically updated across hierarchical levels contribute to individual variability in perceptual inference, supporting models of predictive processing in which both stable priors and adaptive updating mechanisms shape perception over time.

### Multiscale Biases and Decision Dynamics Interact in Shaping Perceptual Inference

Together, these findings provide converging evidence that causality perception reflects a perceptual inference process where priors and decision-making dynamics vary across individuals, but how do these components jointly shape perception? First, our correlational findings suggest that perceptual history contributes to systematic shifts in the starting point of the decision process, supporting the idea that recent perceptual experience and stable internal priors may interact and converge within a unified inferential framework [30]. Indeed, our results indicate that these mechanisms may not operate independently: participants with a greater initial bias toward causality exhibited stronger serial dependence. This relationship highlights how strong priors and heightened sensitivity to perceptual history may interact to reinforce one another, creating a feedback loop in which each causal judgment increases the likelihood of perceiving causality in subsequent events. This perceptual ‘*stickiness’*, potentially driven by underweighted low-level priors and compensatory top-down predictions [20], may create a feedback loop in which biased perceptual decisions become increasingly self-confirming, fostering cognitive rigidity and biased causality attribution as observed in SSD [13,50,51].

We also found that lower decision thresholds were associated with stronger serial dependence, suggesting that participants who commit to perceptual decisions more quickly may be more susceptible to the influence of recent perceptual history. On the other hand, individuals with higher decision thresholds, more typical of ASD-like profiles, showed reduced reliance on perceptual history, consistent with a cautious, sensory-driven processing style requiring greater evidence to update internal representations [27,31].

Furthermore, the point of subjective causality was negatively correlated with both starting bias and drift rate. This suggests that individuals with a stronger initial bias and faster accumulation of evidence in favor of causality required longer temporal delays to perceive non-causal contingencies between visual events, reflecting a more rigid cognitive-perceptual style and potentially explaining higher visual temporal binding thresholds in SSD [21–23].

Overall, our findings support the view that causality perception emerges from the dynamic interplay of predictive stability, perceptual adaptability, and hierarchical precision weighting, revealing systematic individual differences in how perceptual decisions are shaped across the general population.

## Conclusions, Limitations and Future Directions

Causality perception serves as a fundamental mechanism for parsing the sensory world and structuring temporal experience [3]. Understanding its variability thus provides key insight into the architecture of perceptual processing. Our findings reveal that causality perception arises from a hierarchical and dynamic interplay of sensory input, internal priors, and perceptual history, and that these processes vary systematically across trait-based profiles in the general population. Stratifying participants based on ASD- and SSD-like traits revealed behavioral and computational markers of prior- and sensory-driven perceptual styles, highlighting that balanced perceptual processing depends on the flexible integration of these components, a capacity that may be diminished in both SSD and ASD profiles.

Our findings open several avenues for future research to test, replicate, and extend these results. Our trait-based classification relied on self-report measures; future studies incorporating clinical assessments, neuroimaging, and physiological biomarkers could refine and validate these profiles. Moreover, whether similar trait-dependent differences extend to other domains of perceptual inference (e.g., agency, motion prediction, or affective interpretation) remains an open question. Finally, our findings underscore the importance of multidimensional models of predictive processing that account for individual variability across both typical and atypical populations. Future research should further explore how stable priors, dynamic updating, and decision policies interact across sensory and cognitive domains, and how these interactions shape variability in perception, cognition, and behavior.

## Methods

### Participants

A total of 163 volunteers were recruited from the online experiment platform *Prolific* (www.prolific.com). Participants presented normal or corrected-to-normal vision and hearing. Exclusion criteria were self-reported neurological and attention disorders, epilepsy, and photosensitivity. During data collection, the refresh rate of the monitor was recorded for each participant, ensuring a correct timing of stimuli presentation at the desired refresh rate (i.e., 60 Hz). Four participants were excluded from subsequent analyses since they performed the task using a monitor with different refresh rate. We excluded an additional nine participants whose behavioural performance exhibited an Adjusted R² value < 0.9 following logistic fitting (see Data Analysis). The final sample included 150 participants (72 females, mean age=23.7 years, SD=3.79). We emphasised the critical importance of sitting in a dimly lit and quiet room and keeping a viewing distance of ∼50 cm from the screen. All participants gave informed consent by checking the relevant boxes on a consent form *Prolific* webpage and received 9.75$ as compensation. The experiment was conducted in accordance with the Declaration of Helsinki and approved by the New York University Abu Dhabi Human Research Protection Program Internal Review Board (IRB).

### Stimuli and Experimental Design

The task was created with PsychoPy [52] and administered via Pavlovia (https://pavlovia.org/), a web-based platform for the presentation of psychophysics experiments via common web browsers. Visual stimuli were created using Psychtoolbox on MATLAB 2019a (MathWorks, Inc), generating 60 Hz-videos of launching sequence at different time lags. All stimuli were presented on a 60-Hz monitor at a recommended viewing distance of 50 cm against a uniform gray background. Each trial commenced with a fixation cross at the center of the screen presented for 1000 ms, followed by the presentation of a launching sequence. This sequence consisted of a circle moving toward and “colliding with” a second circle, subsequently “causing” the second circle to move in the same direction. The first circle moved toward the second circle for 800 ms before colliding with it. Upon collision, the display remained static, and to manipulate participants’ perception of causality, we introduced a temporal lag at the moment of collision. Specifically, we introduced 10 equally spaced collision lags ranging from 0 to 144 ms, increasing in steps of one frame, to elicit either causal or noncausal impressions. After collision time lag, the second circle moved away for 800 ms before disappearing. Participants were then presented with a question (”*Causal or Noncausal?*”), requiring them to indicate their perception of causality through a forced-choice response as quickly as possible. The stimuli moved vertically either upward or downward on the screen and were consistently presented in the left visual hemifield, with the initial position of the bounced circle aligned with the fixation point. Each combination of sequence direction and collision lag was presented 20 times, resulting in a total of 440 randomized trials divided into three blocks. The entire experiment lasted approximately 30 minutes per participant. Participants were instructed to maintain fixation while attending to the lateralized stimuli and to indicate via key press whether they perceived the interaction between the moving circles as causal. Importantly, they were explicitly advised not to treat the task as a lag detection task but rather to focus on their subjective impression of causality induced by the collision. All participants were naive to the experiment’s purpose.

### Autistic Traits

Autistic traits were assessed using the Autism Spectrum Quotient (AQ; [39]), a self-report questionnaire designed to evaluate multiple aspects of cognitive style, behavioral and socio-communicative patterns, and sensory experiences. The AQ consists of 50 items distributed across five subscales, each comprising 10 items: (i) attention to detail, (ii) attention switching, (iii) imagination, (iv) communication, and (v) social skills. Participants rated each item on a 4-point Likert scale, ranging from “definitely agree” to “definitely disagree,” indicating the extent to which they endorsed each statement. Following the original scoring method (Baron-Cohen et al., 2001), we computed a total AQ score by summing the individual subscale scores, with higher values reflecting greater levels of autistic traits.

### Schizotypal Traits

Schizotypal traits were assessed using the Schizotypal Personality Questionnaire (SPQ; [40]), a self-report measure consisting of 74 items. These items are distributed across nine subscales: ideas of reference, magical thinking, social anxiety, unusual perceptual experiences, constricted affect, lack of close friends, odd behavior, odd speech, and suspiciousness. The subscales are further grouped into three main factors: cognitive-perceptual, interpersonal, and disorganization. Participants responded to each item using a binary format (”Yes” or “No”), indicating whether they endorsed the statement. Following the original scoring method (Raine, 1991), responses were coded as 0 for “No” and 1 for “Yes,” with higher scores reflecting greater schizotypal traits.

### Data Analysis

#### Psychometric Function of Visual Causality Judgments

We fitted a psychometric logistic curve to the percentage of non-causal judgments as a function of collision time lags. We performed the fitting of the psychometric function separately for each participant (mean adjusted R² = 0.956). The adjusted R² of the fitted model was evaluated individually, and participants with poor logistic fitting (adjusted R² < 0.9) were excluded from the statistical analysis (n = 9; final sample = 150). We used a logistic equation and a nonlinear least squares method to fit the proportion of noncausal responses as a function of collision time lags. The formula applied was as follows: *y = 1/(1 + exp(b × (t - x))).* In this equation, *x* represents the time lag, and *y* denotes the proportion of non-causal responses. The lower bound of y was set at 0, and the upper bound was set at 1. The only free parameters of the function were *b* (the slope of the function) and *t* (the 50% threshold), both of which were constrained to assume positive values above zero. We obtained the individual 50% threshold values from the fitting of the psychometric logistic curve, reflecting the point of subjective causality (PSC). Second, to investigate the impact of perceptual history on visual causality judgments, we implemented a serial dependence approach [53], where all trials were divided into two bins based on whether they followed a causal (i.e., t-1 causal) or non-causal (i.e., t-1 non-causal) response trial. We then performed additional logistic fitting of non-causal responses for t-1 causal (mean adjusted R² = 0.923) and t-1 non-causal trials (mean adjusted R² = 0.938), obtaining a PSC estimate for each bin and participant. The difference between the PSC of t-1 causal and t-1 non-causal trials served as a proxy measure of the influence of previous trials on current causality judgments.

Additionally, to further investigate variations in visual causality judgments across each time lag, we examined each participant’s mean reaction times (RTs), separately indexing all trials, t-1 non-causal trials, and t-1 causal trials.

#### Individual differences along the ASD-SSD continuum: Cluster Analysis

To characterize the multiple phenotypes observable within the ASD-SSD continuum, previous studies have employed factorial (e.g., Principal Component Analysis; [14]) and clustering (e.g., k-means/k-median; [42,54) approaches on measures derived from the AQ and SPQ questionnaires. In the present study, we chose to implement a cluster analysis technique rather than a factorial approach, as the latter is based on a variable-centered methodology that tends to underestimate the interindividual variability within a sample [54]. In contrast, cluster analysis is a person-centered approach that reveals commonalities between individuals, thus identifying group-specific relationships between variables that are often obscured in a factorial approach. Accordingly, our sample was stratified using a k-means cluster analysis (Hartigan-Wong method with squared Euclidean distance, and a maximum of 25 iterations to find the optimal clustering solution) based on participants’ z-scored ratings on the five subscales of the AQ (*social skills, attentional switching, attention to detail, communication, and imagination)* and the nine subscales of the SPQ (*ideas of reference, social anxiety, odd beliefs/magical thinking, unusual perceptual experiences, eccentric/odd behavior, no close friends, odd speech, constricted affect, and suspiciousness*). This data-driven approach characterized our sample into three distinct clusters of participants along the ASD-SSD continuum: (i) ASD-like traits participants, (ii) Low Traits participants, and (iii) SSD-like traits participants (see Results section for more details).

#### Hierarchical Drift Diffusion Model (HDDM)

After stratifying our sample into three distinct clusters along the ASD-SSD continuum (see Results section), we implemented a hierarchical drift-diffusion model (HDDM) using Python 3 and the HDDM toolbox [38]. This approach enables the embedding of drift-diffusion models (DDMs) within a hierarchical framework and estimates parameters using Bayesian methods. A key advantage of HDDM over classical DDM is its ability to estimate parameters at both the individual and condition levels simultaneously, based on variable group-level distributions. This reduces the influence of individual outliers and allows for reliable parameter estimation even with a limited amount of data per participant. We employed HDDM to examine whether differences in visual causality judgments across the ASD-SSD continuum could be explained by variations in the starting point of the accumulation process (i.e., initial *bias* in favor of one option; starting point), the speed at which evidence is accumulated (*drift rate*), or the amount of information required to make a decision (*decision boundary*).

Accordingly, we first fitted the HDDM to the RTs distributions for causality responses, separately for each cluster. In our model, we allowed the following parameters to vary as a function of time lag: (i) the drift rate (v), (ii) the separation between decision boundaries (a), and (iii) the starting point of the accumulation process (z). In summary, we estimated the posterior distributions for a total of 3 parameters for each cluster. We initialized HDDM to draw 20,000 posterior samples for each data with the first 5000 samples discarded as burn-in. We estimated the posterior distributions of our parameters by computing four independent Markov Chain Monte Carlo (MCMC). Convergence of the remaining 15000 samples was assessed using visual inspection of the MCMC trace plots values and their autocorrelation to ensure that the models had properly converged. Posterior distributions and MCMC trace diagnostics for all key HDDM parameters (starting point bias, drift rate, and decision boundary) are reported in Supplementary Figure S1, showing well-behaved, unimodal posteriors and stable chain convergence. All parameters exhibited R^ values below 1.01, indicating excellent convergence across sampling chains. To further validate the goodness of fit of our model, we conducted additional HDDM analyses in which the estimated parameters were modelled as varying based on group alone, time lag alone, their interaction, or independently of these factors. We then compared the deviance information criterion (DIC) across models. Additionally, we conducted posterior predictive checks (PPC) to validate the HDDM parameter estimates (see: *Supplementary Materials S2 and S3*). Specifically, we used the posterior distribution of the model parameters to generate multiple simulated datasets, replicating response times and choices for each trial, participant, and experimental condition. We then assessed the model’s goodness-of-fit through visual inspection of the simulated data against the observed data, thus ensuring the model’s validity.

#### Statistical Analysis

First, we performed a one-way ANOVA on the PSC to test whether visual causality judgments were influenced by cluster (between-subjects factor with three levels: ASD-like, Low Traits, SSD-like). Furthermore, to investigate potential differences across clusters in the influence of perceptual history (i.e., serial dependence; trial n-1) on visual causality judgments, we performed 2 × 3 repeated measures ANOVA (rmANOVA) on the PSC separately computed for trials following causal (t-1 causal) and non-causal (t-1 non-causal) judgments, with t-1 judgment (two levels: t-1 causal, t-1 non-causal) as the within-subjects factor and cluster (three levels: ASD-like, Low Traits, SSD-like) as the between-subjects factor. Additionally, to assess the extent to which prior perceptual history influenced current causality judgments across clusters, we computed the PSC difference between *t-1* non-causal and *t-1* causal trials, with larger differences indicating a stronger serial dependence effect. We then conducted a one-way ANOVA on the resulting PSC difference index, with cluster as the between-subjects factor (three levels: ASD-like, Low Traits, SSD-like).

Next, we conducted a 10 × 3 repeated-measures analysis of variance (rmANOVA) on RTs observed across all trials to examine whether the speed of causality judgments was influenced by time lags (within-subject factor: ten levels) and cluster (between-subjects factor: three levels). Additionally, to investigate potential differences across clusters in the influence of perceptual history on RTs, we performed a 10 × 2 × 2 rmANOVA, with time lags and t-1 causal vs. t-1 non-causal trials as within-subject factors, and cluster as the between-subjects factor. The Greenhouse–Geisser correction was applied where necessary to account for violations of the sphericity assumption. All post-hoc p-values were Holm-corrected.

Regarding HDDM, we characterized the posterior distribution of each parameter (i.e., Bias, Drift Rate, Decision Boundary) using its median value and high-density interval (HDI), separately for each cluster. For statistical inference, we assessed between-groups differences by comparing posterior distributions by subtracting their HDIs (Hennig et al., 2025). Specifically, for each contrast of interest, we computed 15,000 differences based on the 15,000 iterations of each parameter estimation, and the resulting significance threshold was used to determine the boundaries of the high-density interval (HDI) distribution. If the HDI distribution of mean differences did not include zero, the condition effect was considered credible and thus statistically significant [45].

Statistical analyses were conducted using JASP (version 0.17.2.1; [55]), Python 3 and MATLAB (version R2021b; The MathWorks Inc., Natick, MA, USA).

## Acknowledgements

This work was supported by the NYUAD Center for Brain and Health, funded by Tamkeen under NYU Abu Dhabi Research Institute grant CG012.

## Author contributions

Conceptualization, G.M. D.M.; Methodology G.M. D.M.; Formal Analysis, G.M.; Software, G.M.; Visualization, G.M.; Funding Acquisition, D.M.; Writing - Original Draft Preparation, G.M. D.M.; Writing – Review & Editing, G.M. D.M.; Supervision, D.M.

## Data availability

Data are shared open-access at https://osf.io/6cmna/.

