## Supplementary Materials for "Atypical weighting of sensory evidence and priors in causality perception along the autism–schizotypy continuum"

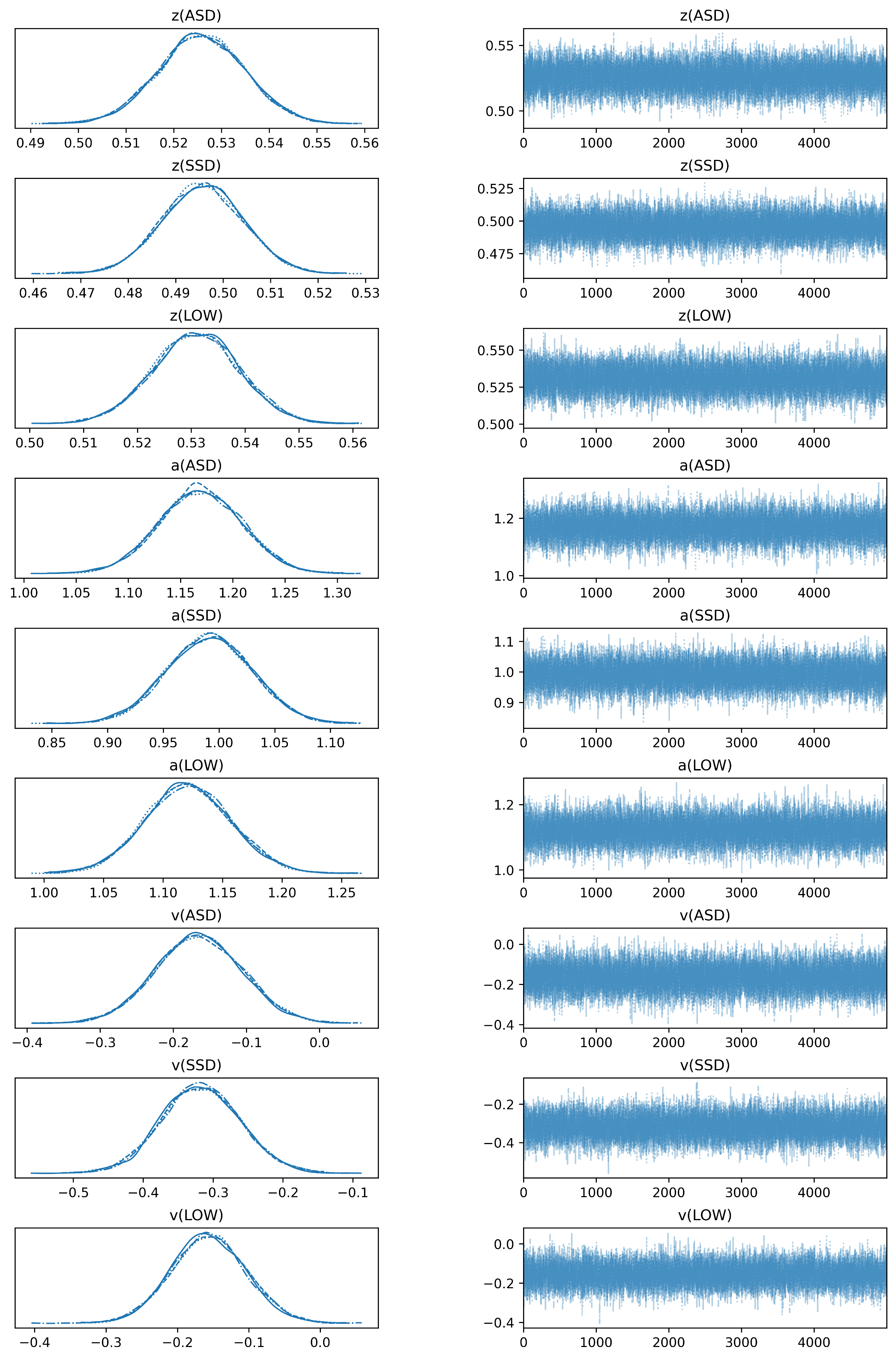

**Figure S1.** **Traceplots and Posterior Distributions for each HDDM parameter.** Posterior distributions (left column) and Markov Chain Monte Carlo (MCMC) trace plots (right column) for the key decision-making parameters estimated via the Hierarchical Drift Diffusion Model (HDDM), shown separately for each trait group: ASD-like, SSD-like, and Low Traits. The posterior distributions (left) display well-formed, unimodal, and symmetric shapes, suggesting stable and reliable parameter estimation across groups. The trace plots (right) show the evolution of posterior samples across MCMC iterations, with consistent mixing and no visible drift, indicating good convergence of the model chains.

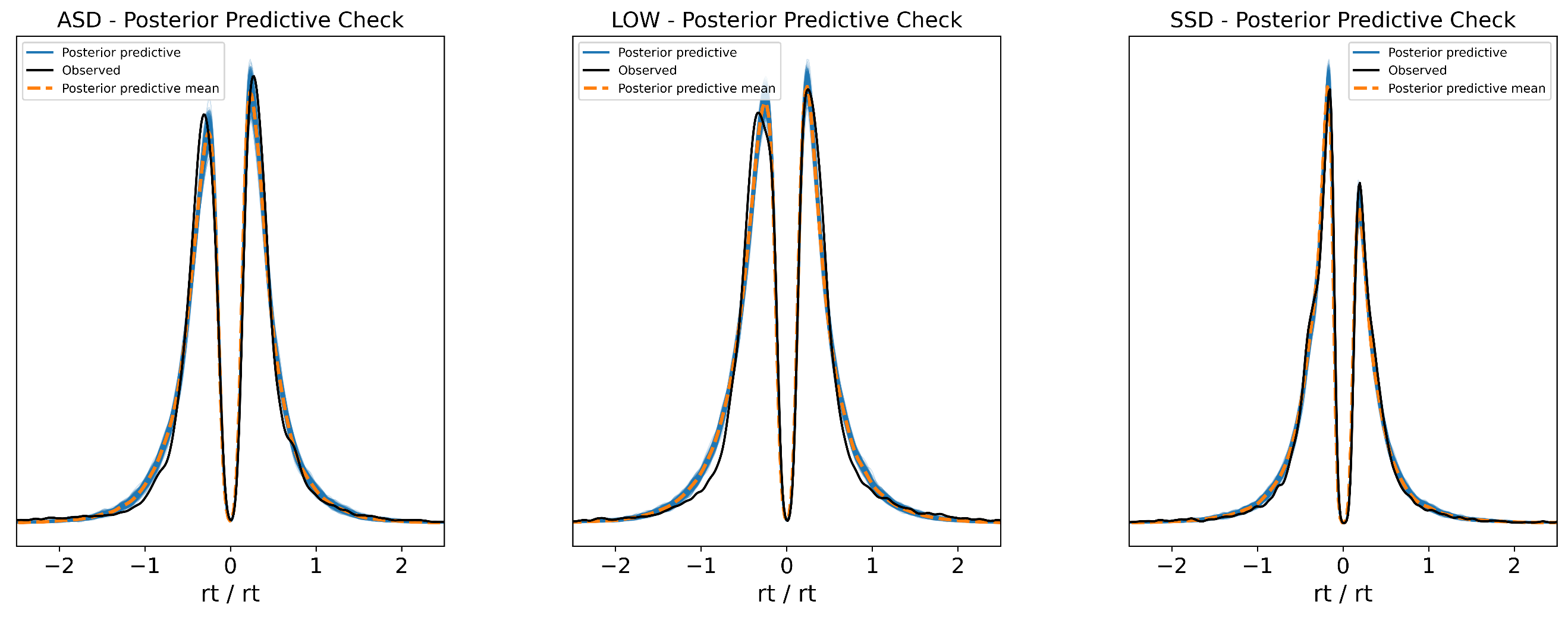

**Figure S2.** **Group-level** **Posterior predictive check (PPC) for the HDDM (Hierarchical Drift Diffusion Model), separately for the ASD-like, Low Traits, and SSD-like participant groups.** Each panel shows the observed reaction time (RT) distribution for each group (black line), overlaid with posterior predictive samples (blue shaded area) and the posterior predictive mean (orange dashed line) generated from the model. The close alignment between the observed data and the posterior predictive distributions indicates that the model accurately captures the RT distributions across all three groups. Both the shape and central tendencies of the distributions are well recovered, suggesting a good overall model fit. This supports the validity of the estimated decision parameters (bias, drift rate, and boundary separation) at the group level and provides evidence that the HDDM reliably generalizes across distinct cognitive-perceptual profiles.

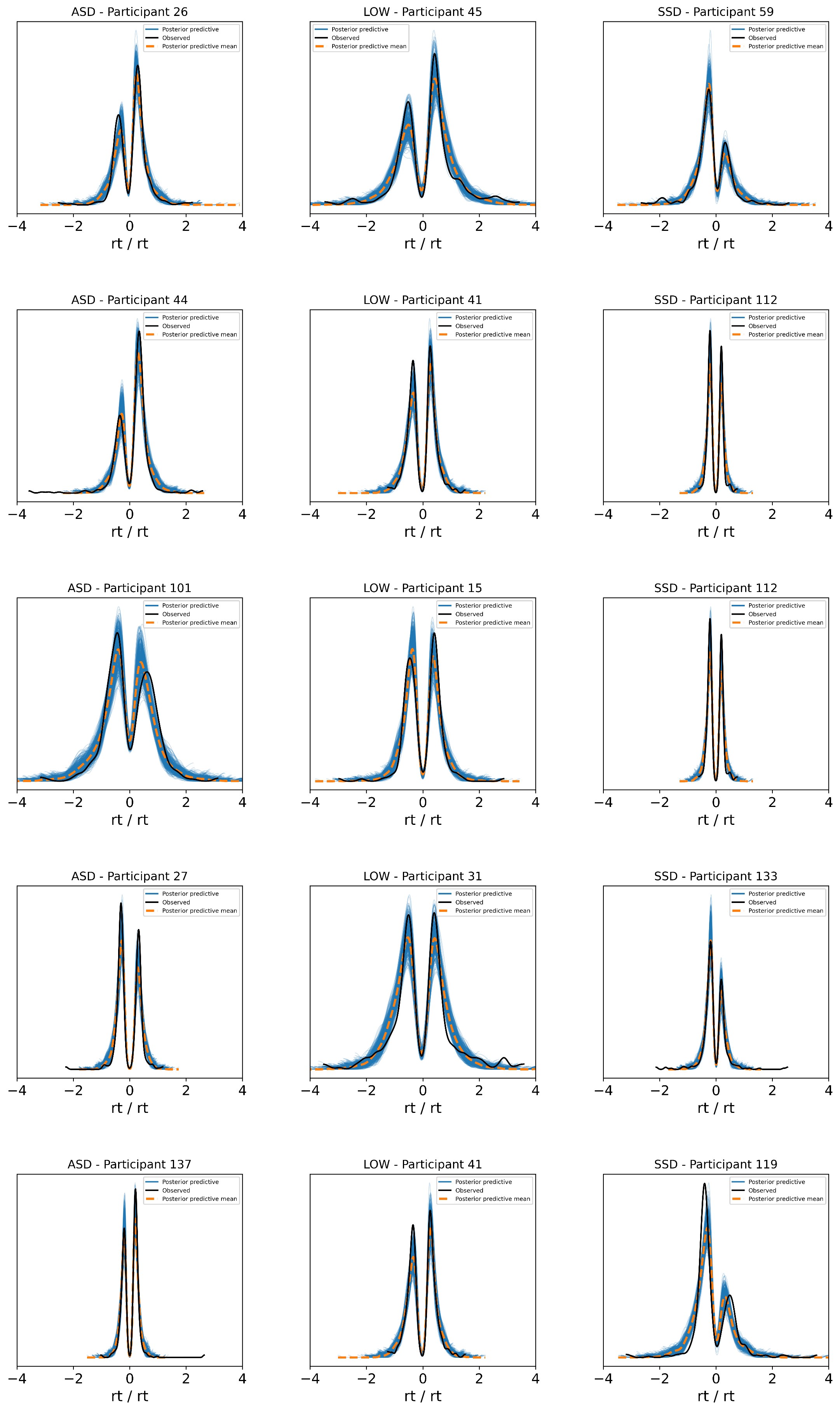

**Figure S3.** **Individual-level Posterior predictive check (PPC) for the HDDM (Hierarchical Drift Diffusion Model)**. Posterior predictive checks at the individual level for randomly selected participants from the ASD-like, Low Traits, and SSD-like groups. Each subplot displays the observed reaction time (RT) distribution for a single participant (black line), the corresponding posterior predictive mean (orange dashed line), and the posterior predictive distribution samples (blue shaded area) simulated from the fitted HDDM. Across individuals, the observed data are consistently well captured by the model-generated predictions. The HDDM successfully reproduces both the overall shape and variance of the empirical RT distributions, including the characteristic bimodality resulting from fast and slow responses. The tight overlap across participants confirms that the model not only fits group-level trends but also robustly accounts for individual variability in decision-making dynamics.
